# Wearable MEG data recorded during human stepping

**DOI:** 10.1101/2025.02.11.637004

**Authors:** Meaghan E. Spedden, George C. O’Neill, Timothy O. West, Tim M. Tierney, Stephanie Mellor, Nicholas A. Alexander, Robert Seymour, Jesper Lundbye-Jensen, Jens Bo Nielsen, Simon F. Farmer, Sven Bestmann, Gareth R. Barnes

## Abstract

Non-invasive spatiotemporal imaging of brain activity during large-scale, whole body movement is a significant methodological challenge for the field of movement neuroscience. Here, we present a dataset recorded using a new imaging modality – optically-pumped magnetoencephalography (OP-MEG) – to record brain activity during human stepping. Participants (*n*=3) performed a visually guided stepping task requiring precise foot placement while dual-axis and triaxial OP-MEG and leg muscle activity (electromyography, EMG) were recorded. The dataset also includes a structural MRI for each participant and foot kinematics. We validate the fidelity of the OPM data by showing movement-related beta band desynchronization source localised to the primary motor cortex. This multimodal dataset offers a resource for methodological development and testing for OPM data (e.g., movement-related interference rejection), within-subject analyses, and exploratory analyses to generate hypotheses for further work on the neural control of human stepping.

## 1. Background

Walking is a fundamental behavior integral to all humans and animals that enables us to move around our environment. Evidence indicates that, similar to other animals, humans have a spinal cord network, known as a central pattern generator, that is responsible for generating the fundamental walking pattern^1,2^. However, in humans, the function of this network is more reliant on input from the brain and has been adapted to meet the specific demands of bipedal walking^3^.

Studying brain activity non-invasively during large-scale movements (like walking) is methodologically challenging due to the complex, multi-limb, multi-joint actions, and whole-body translation involved. Consequently, imaging methods requiring participants to remain still (e.g., functional magnetic resonance imaging, fMRI, and cryogenic magnetoencephalography, MEG) can use simplified models based on imagined movement or isolated limb movement. In this context, electroencephalography (EEG) studies have contributed significantly to mapping the spectral and temporal features of the cortical network involved in the control of human walking^4–9^ as well as characterizing the cortical influence on leg muscle activity during walking^10–12^. These insights have played a role in refuting the traditional view that the human cortex plays a limited role in stereotyped walking.

A new approach that holds significant potential for further progress in this field is optically-pumped magnetoencephalography (OP-MEG)^13–15^. OPMs are magnetic sensors that do not require cryogenic cooling, which means that they can be positioned within a few millimeters of the scalp in wearable arrays, offering a flexibility similar to EEG^16,17^. Magnetic field-based imaging also offers the advantage of improved spatial resolution relative to EEG as the measured fields are minimally distorted by differences in tissue conductivity^18,19^. MEG also offers lower sensitivity to muscle artifacts relative to EEG^20^, which is critical for paradigms involving head movement.

Using OPMs to record brain activity during large scale movement is however not without significant challenges^16,21^. Sensor movement through the ambient magnetic field generates large, low-frequency artifacts that can exceed both the magnitude of neural activity as well as the sensor dynamic range, making the data unusable. Importantly, recent developments in both OPM technology and analytical tools^16,17,22–24^ mean that we can begin to shift our perspective from considering movement as a confound to studying the neurophysiology of large-scale, natural movement.

In this data resource paper, we present an OP-MEG dataset from three participants performing a visually guided stepping task. The task captures a fundamental aspect of walking behavior, the ability to precisely place the feet based on visual input^25^. Walking is modelled by discrete steps in this task for practical considerations (cabling, stimulus presentation), but it is important to note that it is certainly possible to perform continuous walking with the existing OPM systems^17^.

The dataset is multimodal, including OP-MEG, structural MRI, foot kinematics, and leg muscle activity (electromyography, EMG) from each participant. We have used this dataset to establish proof-of-principle that OP-MEG can record high fidelity brain activity during large scale, whole body movement. We present this work in the present paper as technical validation, demonstrating that we can image movement-related modulations of beta band activity^26,27^ in the sensorimotor cortex using OP-MEG.

To the best of our knowledge, this is the first publicly available ambulatory MEG dataset. It allows interested researchers to explore the potential of OP-MEG, providing a basis for generating hypotheses about the spatiotemporal dynamics of stepping from an imaging perspective. For example, the OP-MEG data and individual MRIs make it possible to perform high fidelity source reconstruction analysis to identify brain networks that are active during step planning and execution.

The inclusion of EMG data also means that the spatial features of functional connectivity with muscle activity can be explored using functional and directed connectivity metrics like cortico-muscular coherence^12^. Task performance data and kinematics are also included in the dataset, allowing the neural activity patterns to be linked to movement features and task performance. Finally, an EEG study has been published using the same stepping paradigm^7^, which allows the comparison of EEG results to OP-MEG results. Given that the dataset only comprises data from three participants, it is primarily suited for within-participant, hypothesis generating analysis as a foundation for larger studies.

We also suggest that the dataset can be used to evaluate, compare, and develop approaches to movement-related interference rejection in OPM data. We use current state-of-the art spatial filtering approaches^23,28^ in our technical validation that permit the imaging of movement-related modulations of beta band activity, however, OPM analysis is still in its early stages and will benefit continued refinement and innovation.

## 2. Methods

### 2.1 Participants

Three healthy participants (Age 55, 33, and 30; all male) participated in this study. Written, informed consent was obtained prior to participation and the experimental protocol was approved by the University College London Research Ethics Committee.

### 2.2 Stepping task

Participants performed a visually guided stepping task while we recorded OP-MEG and concurrent EMG from the tibialis anterior (TA) muscle.

The stepping task was an adapted version of a visually guided walking paradigm used in previous work^7,12,29^ which demonstrated the presence of cortico-muscular and cortico-cortical coupling during stepping.

The task required participants to take single steps using their right leg, aiming for virtual stepping targets. The stepping target and real-time position of the stepping leg were projected on a screen, represented as a magenta square and blue circle, respectively (Figure 1A). The task involved using the real-time visual feedback of the stepping leg position relative to the stepping target to adjust step length and hit the target. Stepping leg position was tracked using a set of six infrared cameras at 120 Hz (Optitrack, Flex 3, Natural Point, Inc.) to record the position of retro-reflective markers placed on the stepping foot. The markers were used to construct a rigid body of the foot, and its coordinates were streamed using the Motive (Natural Point, Inc.) NatNet SDK to MATLAB, where a custom script ran the stimulus presentation.

**Figure 1.**
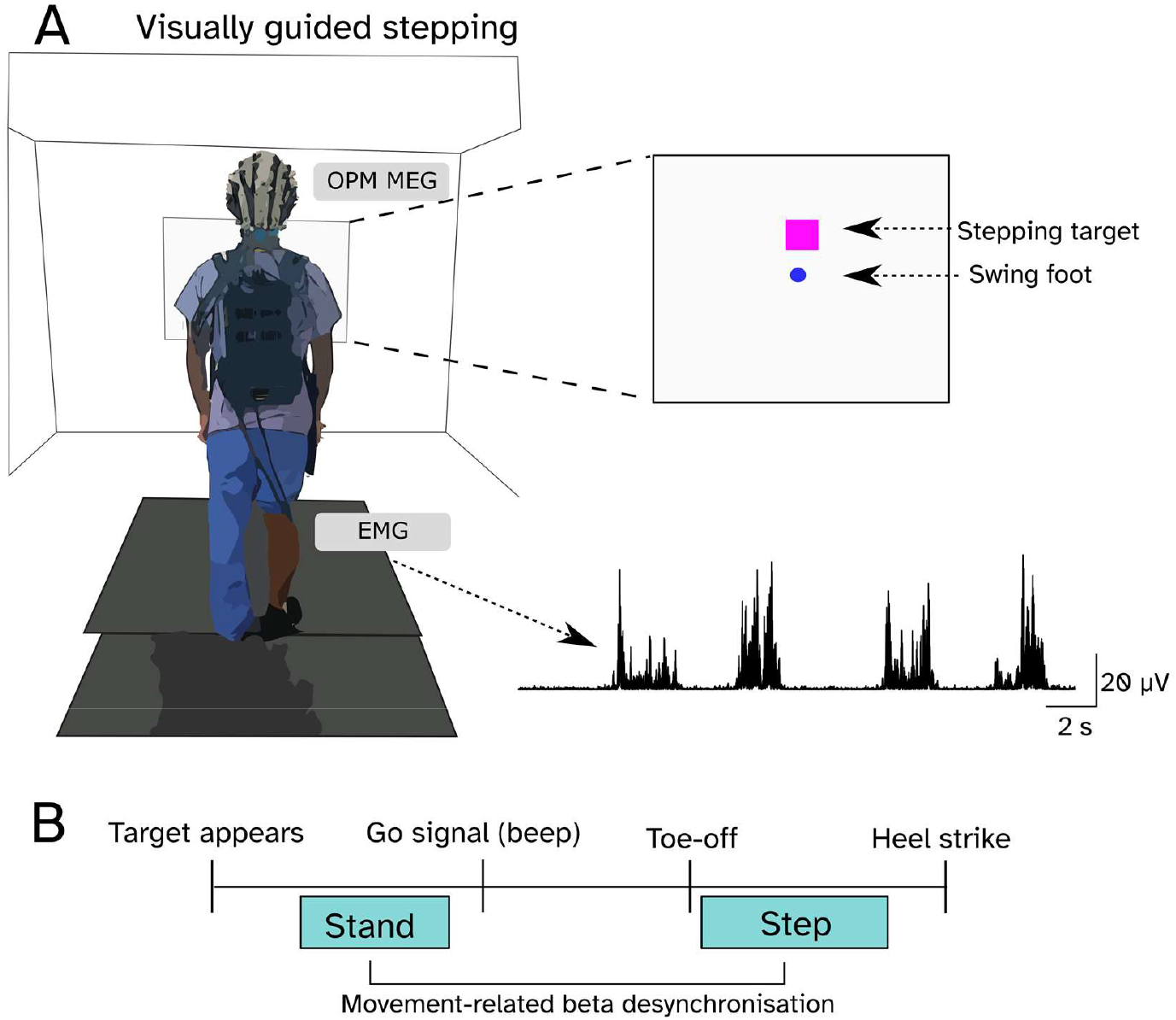
Experimental setup. In the visually guided stepping task (A) participants took single steps forwards aiming to hit a virtual target (magenta square) with a blue circle reflecting the position of their right foot. OP-MEG and EMG from the right tibialis anterior (TA) were recorded.

Prior to the start of the experiment, the virtual target distance was adjusted to represent a comfortable step length for the participant (preferred step length). During recordings, target distance (step length in the anterior–posterior direction) was drawn randomly from 3 possible values: preferred step length, preferred length +5 cm and preferred length -5 cm. Participants began each trial standing quietly and took a step forward with the right leg aiming to hit the magenta square (target) with the blue circle, which moved with the right foot. The trial started when the target was projected on the screen, and participants were instructed to initiate the step when they heard a beep serving as the go signal. The trial was completed when the left foot was placed next to the right foot, and the participant then returned to the starting position, which was marked as an open circle on the screen. Trial duration was ∼10 seconds, and 5-6 blocks of 30 steps each (5 for participants 1 and 2; 6 for participant 3) were recorded.

The custom MATLAB script sent synchronizing triggers to OPM and EMG acquisition systems upon target appearance for each trial. The script also wrote rigid body kinematics, trigger timing, target position, 2D foot position projection (blue circle) to a text file.

### 2.3 Electrophysiological recordings

#### 2.3.1 Optically Pumped Magnetoencephalography (OP-MEG)

The experiments were performed in an MSR (Magnetic Shields, Ltd., Staplehurst, UK; internal dimensions 3 x 4 x 2.2m). The room was degaussed prior to the start of the experiment. Dual axis and triaxial OP-MEG sensors (QuSpin Inc., Louisville, CO, USA) were positioned in sockets in a rigid scanner-cast constructed from each participant’s structural MRI (Participant 1: 30 dual-axis sensors; participant 2: 27 dual-axis sensors; and participant 3: 47 triaxial sensors. See Figure 2 for sensor layouts). This ensures accurate co-registration, maximal signal for any head size, and the rigidity minimizes sensor and cable movement relative to the head. For participants 1 and 2, OP-MEG data was acquired using a National Instruments acquisition system and a custom LABVIEW program with a sampling frequency of 6000 Hz. An antialiasing 500 Hz low-pass filter (60th order FIR filter combined with a Kaiser window) was applied before data were down-sampled offline to 2 kHz. Sensors were operated in a mode with a dynamic range of ± 4.5 nT. For participant 3, we used the Neuro-1 acquisition system (QuSpin Inc., Louisville, CO, USA) consisting of exclusively tri-axial sensors (in open loop mode) with a sampling frequency of 375 Hz. Both acquisition systems have an intrinsic bandwidth of 0-135 Hz (due to the properties of the vapour cell). In the Neuro-1 acquisition system the manufacturer has additionally implemented a high order digital low pass FIR filter at 150 Hz.

**Figure 2.**
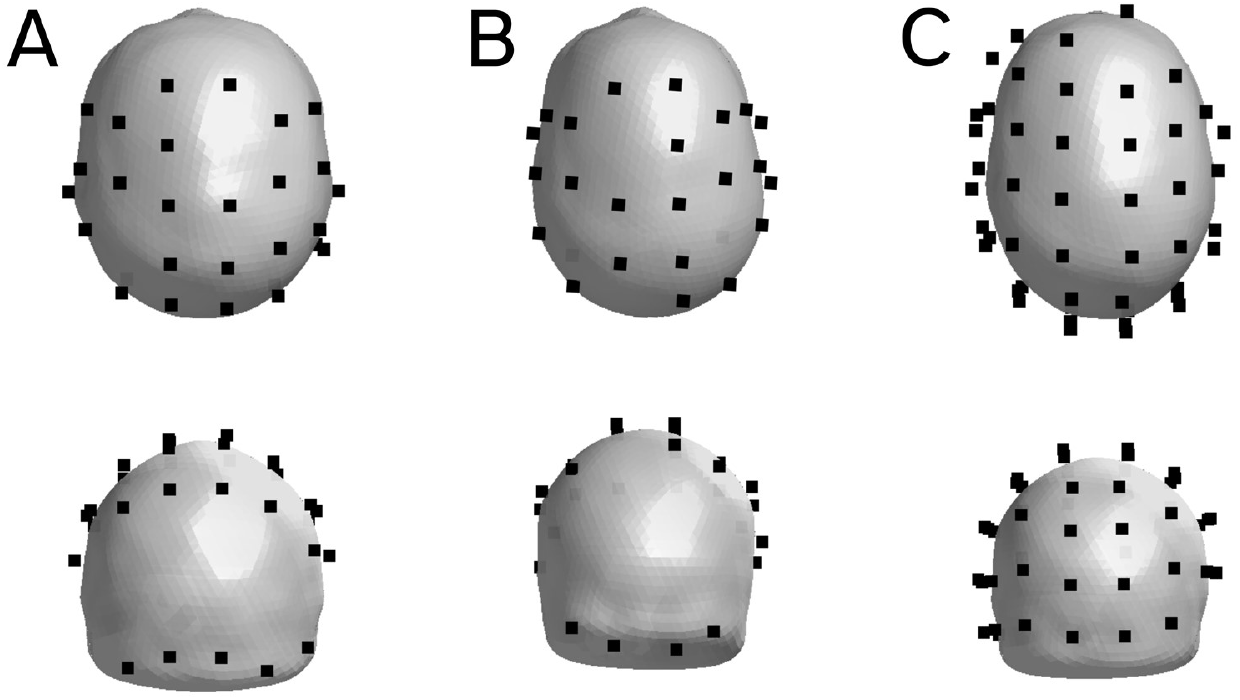
OPM sensor coverage. Coverage is shown for each participant (A-C, respectively). Sensors are depicted as black squares on the scalp mesh derived from each participant’s MRI. First row: superior view. Second row: posterior view.

#### 2.3.2 EMG

EMG was recorded from the right tibialis anterior (TA) muscle. Two surface electrodes (2 cm diameter, Natus Neurology, Inc.) were placed over the centre of the muscle (ca. 2 cm between electrodes) and a ground electrode was positioned on the right lateral malleolus. To prevent EMG data collection from interfering with OPM signals, we passed EMG cables through waveguides so that EMG signals were amplified, filtered, and digitized outside the MSR (Amplification: x 1000; hardware filtering 3 to 100 Hz and 50 Hz notch D-360 amplifier; digitization at 1000 Hz, and 1401 data acquisition unit, Cambridge Electronic Design, UK). Signals were recorded using Spike2 software (v10.05).

### 2.4 Analysis

The technical validation analysis was performed in MATLAB (R2021b) and code is available on GitHub (https://github.com/meaghanspedden/stepping_opm_data). We used the development version of Statistical Parametric Mapping (SPM; https://github.com/spm)^30^ to perform source imaging of movement-related beta power.

#### 2.4.1 Preprocessing

OPM data were imported into SPM and resampled to 1000 Hz. OPM power spectra were then visually inspected for bad channels (i.e., large deviations from median power and/or manufacturer’s noise floor). Harmonic models, i.e., homogenous field correction (HFC) or adaptive multipole models (AMM), were applied to OPM data for interference suppression^23,28^ depending on the number of OPM channels. HFC was applied to data from participant 1 and 2 (number of channels < 120), whereas AMM was applied to data from participant 3 (number of channels > 120)^23^. HFC models the interference as a spatially constant field and can substantially reduce environmental noise over a broad range of frequencies^21^. AMM makes not only a model of the external interference, but also the fields arising within the head and a third partition of signal that does not conform to either model; this adds further immunity against magnetic sources of interference^23^.

OPM and EMG data were high pass filtered at 5 and 10 Hz, respectively, then low pass filtered at 45 Hz, and finally notch filtered at 49-51 Hz to remove residual line noise. Initially, OPM and EMG data were epoched into single trials (i.e., each step) based on triggers indicating stepping target appearance. The standing period was defined as -1.4 to -1.9 s before target presentation for all participants. EMG signals in this interval were visually inspected to confirm that participants were standing still. The stepping period was defined individually for each participant based on the timing of the first EMG burst indicating the swing phase of the step (visual inspection). This period was from 1.5 to 2 s after the go signal for participant 1, 1.6 to 2.1 for participant 2, and from 2 to 2.5 seconds for participant 3. The 500-ms duration was chosen to avoid artifacts related to the heel strike during the stepping period^7^. Pre-processed data were visually inspected and any remaining outlier trials where channels exhibited large deflections were removed.

#### 2.4.2 Source imaging

We used the DAiSS toolbox for SPM to perform source analysis of OPM data during stepping compared to standing using beamforming^31^. Beamformer weights were constructed for 500 ms duration in both standing and stepping time windows, i.e., a common filter, for the beta band (15-30 Hz). The source space comprised a 3D grid covering the whole brain volume (bounded by the inner skull) with 10 mm resolution and a single shell volume conductor^32^. Sensor locations and orientations were innately in MRI space because each scanner cast was constructed from the participant’s MRI. Data were transformed to MNI space in SPM^30^ for consistency and ease of interpretation. The transform was achieved with an affine transformation between fiducials in native and MNI space. Source orientation was optimized for maximal power^33^. A volumetric beta power image was printed for each trial and condition, and standing and stepping were compared (within-participant) using a paired t-test in SPM with the contrast standing > stepping. Note that stepping data are pooled over different step lengths (preferred step length ±5 cm. The resulting images were thresholded using Random Field Theory correction with a significance level of p < 0.05.

#### 2.4.3 Kinematics, EMG, and task performance

We also evaluated the validity of the kinematic and EMG data using visual inspection of position curves and rectified, smoothed EMG signals. Kinematic data were irregularly sampled so first were interpolated to 100 Hz then low pass filtered at 5 Hz. EMG data were rectified by taking the absolute value of the signal, making all values positive, then smoothed with a moving average (window size 100 samples Task performance was calculated as absolute error in the anterior-posterior direction between step endpoint and the centre of the target.

Supplementary Table 1 presents an overview of data files and their contents.

## 3. Results

### 3.1 OPM data

We established the validity of our OPM data by demonstrating movement-related modulations of beta band power during stepping relative to standing. Beta band activity exhibits well-characterized modulations time-locked to planning, initiation, and execution of movement^34^ and is a robust effect^26,27^ occurring in the sensorimotor cortex.

Significant beta band desynchronization was observed in the sensorimotor cortex across all three participants (Figure 3). For all three participants desynchronization was observed bilaterally. Maximal t-statistics were localized to MNI -34 4 70 mm for participant 1; 18 -10 68 for participant 2; and -40 -18 44 for participant 3.

**Figure 3.**
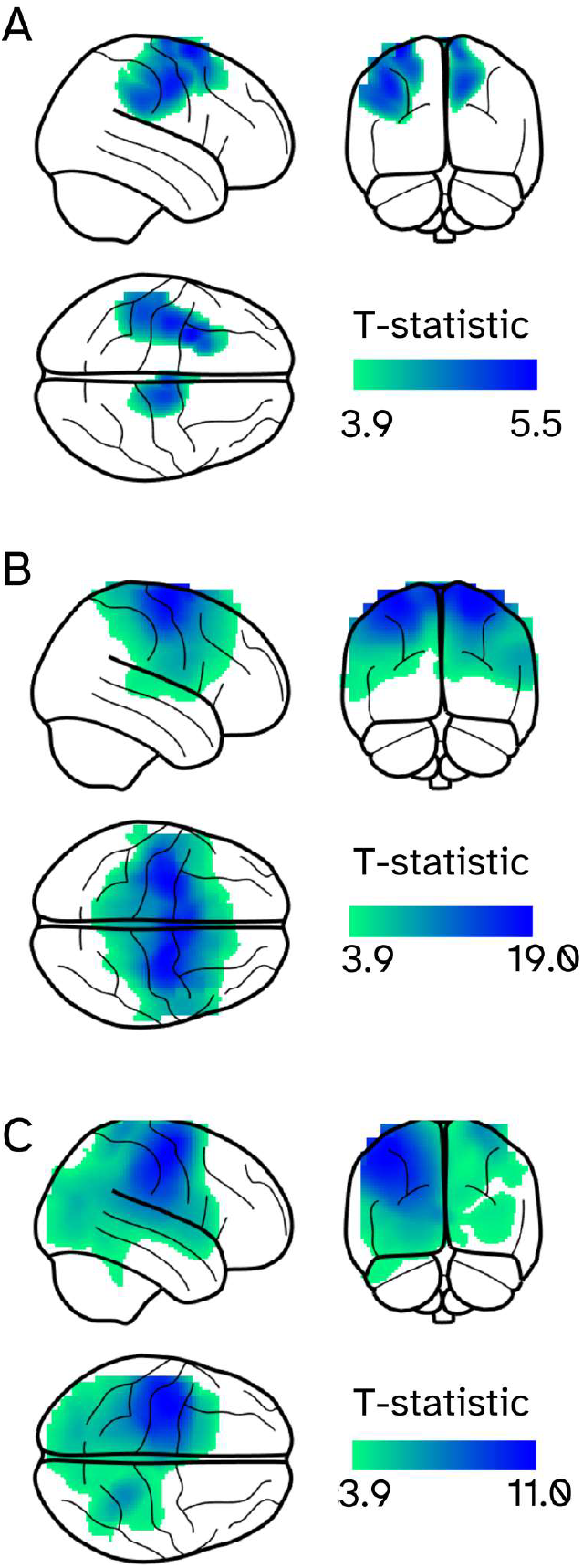
Movement related beta band modulations. Thresholded (p < 0.05, FWE corrected) contrast of stepping versus standing ([1 -1]) in the beta band (15-30 Hz), quantified as a T-statistic for participants 1, 2, and 3, respectively (A-C). Stepping data are pooled over different step lengths.

### 3.2 Kinematics, EMG, and task performance

The timing of the EMG and kinematic signals corresponded well and reflected the expected trajectory and muscle activity patterns of stepping (Figure 4)^35^. This alignment reflects the coordination between neural activation and mechanical output. The EMG patterns exhibited double peaks, which is characteristic of TA muscle activity during walking^12^.

**Figure 4.**
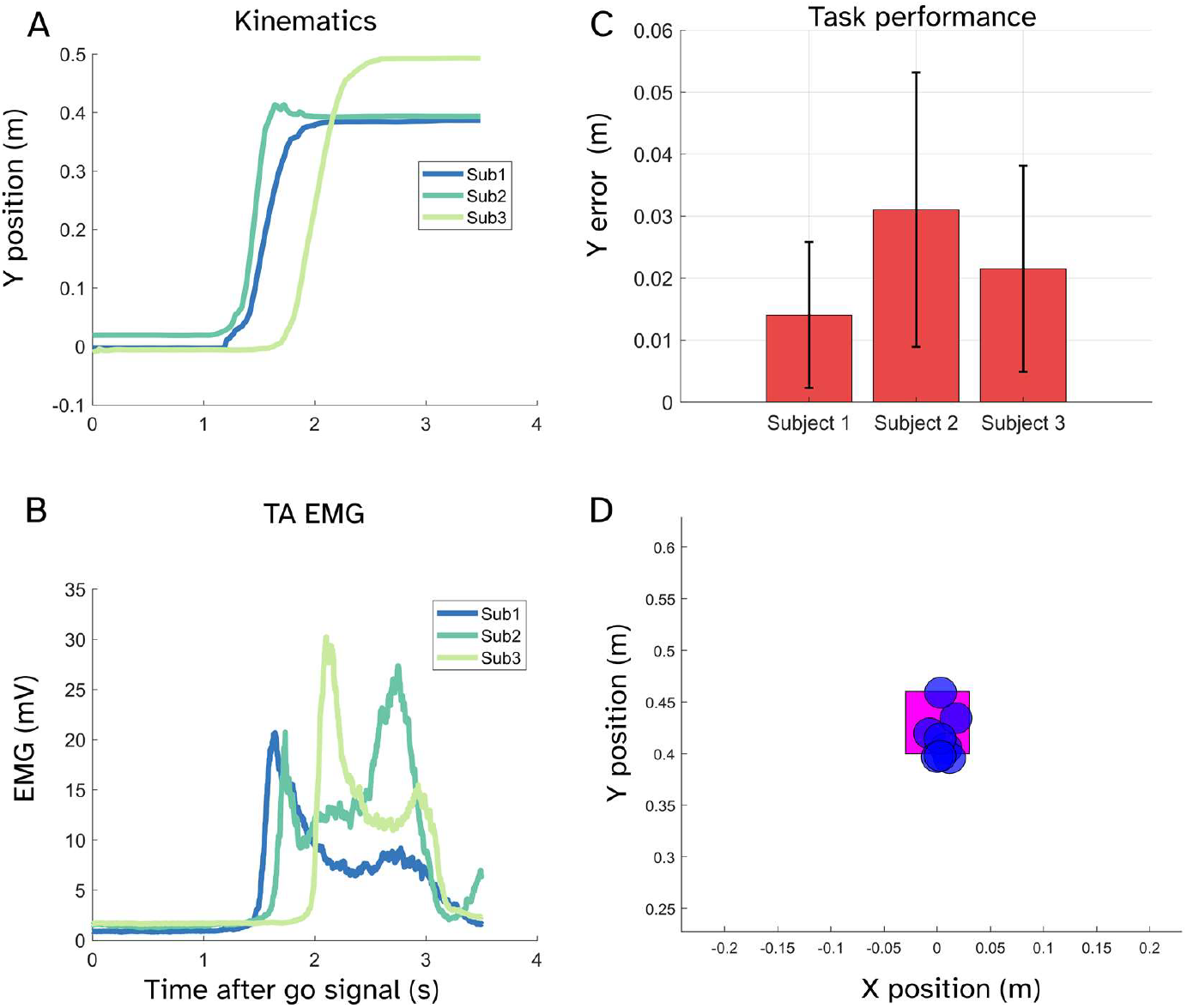
Kinematics, EMG, and task performance. Median foot position across steps (for one of the three target positions) for each participant (A). Median rectified, smoothed EMG trace for each participant (B). Task performance quantified as mean and standard deviation for error in the anterior-posterior direction for each participant (C). Example of foot endpoints relative to the stepping target (D).

## 4. Discussion

We found that all three participants exhibited significant beta band desynchronization in the sensorimotor cortex. These results align with existing literature demonstrating the robust and well-described phenomenon of beta band event-related desynchronization in the sensorimotor cortex during movement compared to rest^36^. The observed bilateral desynchronization is likely attributable to the nature of the whole-body movement, where the stance leg is pushing off while the other swings forward.

This dataset provides the first of its kind resource for exploring the spatiotemporal dynamics of stepping using OP-MEG. Individual MRIs enable high-fidelity source reconstruction to identify brain networks involved in step planning and execution, while EMG data supports functional connectivity analyses using metrics like cortico-muscular coherence. Task performance and kinematic data further allow for linking neural activity patterns to movement features and behaviour.

Although limited to three participants, the dataset is ideal for within-participant analyses and comparisons with EEG data collected using the same paradigm^7^. It also serves as a tool for refining movement-related interference rejection methods.

Another potential limitation is that these data were recorded across two different OPM systems. For instance, the Neuro-1 system has more channels, which can improve spatial filtering and denoising^23^, and its electronics produce less noise, providing more accurate timing, especially for frequencies above 100 Hz. The potential for cable noise and system-generated interference, particularly from the electronics, may affect data from certain brain areas, but with appropriate preprocessing (e.g., using spatial filters like AMM or beamformers), the data are comparable. These differences are more relevant for higher-frequency analyses or regions like the occipital cortex or cerebellum, but for our focus on motor-related brain activity, the signal remains comparable across participants, with the primary concern being slight differences in the analysis due to the varying noise profiles. A comparison of the electronics can be found in ref^37^ . These limitations aside, it is encouraging that the results presented are robust within subjects and across measurement systems.

Software packages suitable for analysing the data include SPM^38^, Fieldtrip^39^, and MNE-Python^40^ and the documentation for each of the three packages contains OPM analysis tutorials. OPM data recorded during stepping is noisy, so we recommend careful interference rejection in preprocessing of this data. We refer to the code used for technical validation in the present work and recent papers on OPM interference rejection^21,23,28^ as guidance.

## 5. Conclusion

By combining OP-MEG with EMG, structural MRI, and foot kinematics, this dataset offers a comprehensive tool for methodological development and hypothesis generation in the field of movement neuroscience.

## Code Availability

The MATLAB code for our validation analysis is available on GitHub (https://github.com/meaghanspedden/stepping_opm_data).

## Data Availability

The data can be found on Mendeley Data with doi 10.17632/p3dfxmky46.2.

## Author contributions

M.E.S, J.L.J., J.B.N., S.F.F., and G.R.B. devised the project. G.C.O, R.C.T, T.W., S.M., T.T., N.A., R.S., M.E.S., and G.R.B. wrote software. All authors contributed to the writing and editing of the manuscript.

