## Supplementary material for "Wearable MEG data recorded during human stepping"

### 1. Data Records

Each dataset comprises OP-MEG data; EMG data; structural MRI; and kinematics/task performance information. All data are de-identified and available on Mendeley Data with DOI:  10.17632/p3dfxmky46.2, and data are predominantly organised according to Brain Imaging Data Structure (BIDS). BIDS is a standard for organizing and describing neuroimaging and behavioural data to facilitate sharing, analysis, and reproducibility. BIDS provides a consistent framework for naming files, defining metadata, and structuring datasets, making it easier for researchers to collaborate and integrate data from different studies. While this does introduce some redundancy, such as using a 'ses' (session) folder when there is only one session per participant, it is important for ensuring compatibility with other datasets. Note that BIDS does not yet officially support OPMs, so we have used traditional MEG conventions. An overview of files and their contents are presented in Supplementary Table 1. Note that we include a README.txt file in the root repository detailing synchronising trigger information.

| Filename template | Contains | Notes |
| --- | --- | --- |
| sub-OP00XXX_ses-001_task-**stepping**_run-XXX_meg.bin  sub-OP00XXX_ses-001_task-**noise**_run-XXX_meg.bin | Raw OPM data | Each dataset also includes a noise (empty room) recording (task-noise). |
| sub-OP00XXX_ses-001_task-stepping_run-XXX_meg.json | Meta data for OPM |  |
| sub-OP00XXX_ses-001_task-stepping_run-XXX_channels.tsv | OPM channel name and type | For subject 00159 channel 39’s name is formatted differently because we did not record from this channel. This does not affect processing but needs to remain in tsv file for metadata compliance. |
| sub-OP00XXX_ses-001_task-stepping_run-XXX_positions.tsv | OPM sensor positions and orientations | Channel name, position, and orientation.  OPM sensors in same coordinate space as MRI |
| sub-OP0XXX_ses-001_task-stepping_run-XXX_emg.tsv | EMG data | EMG signal from right anterior tibial muscle |
| sub-OP0XXX_ses-001_task-stepping_run-XXX_emg.json | Meta data for EMG |  |
| OP00XXX-defaced.nii | Structural MRI |  |
| sub-XXX_ses-001_task-stepping_recording-kinematics_beh.tsv | Kinematics and task information | 3D foot (rigid body) position; 2D foot and target position; trigger timing |

***Supplementary table 1.*** *Overview of file name organisation and content.*
